# Blimp-1 in adipose resident Tregs controls adipocyte beiging and obesity

**DOI:** 10.1101/2019.12.14.874693

**Authors:** Lisa Y. Beppu, Xiaoyao Qu, Giovanni J. Marrero, Allen N. Fooks, Adolfo B. Frias, Katherine E. Helfrich, Ian Sipula, Bingxian Xie, Simon C. Watkins, Amanda C. Poholek, Michael J. Jurczak, Louise M. D’Cruz

## Abstract

Crosstalk between the immune system and adipocytes is critical for maintaining tissue homeostasis and regulating chronic systemic inflammation during diet-induced obesity (DIO). How visceral adipose tissue resident regulatory T cells (aTregs) signal to adipocytes in the visceral adipose tissue (VAT) is not understood. Here we show that Treg-specific ablation of the transcriptional regulator Blimp-1 resulted in increased insulin sensitivity, decreased body weight and increased Ucp-1 in adipocytes in high fat diet (HFD)-fed mice. Mechanistically, we demonstrate that Blimp-1 drives IL-10 production in Tregs, thus suppressing beiging and energy expenditure in adipocytes. Moreover, IL-10 mRNA expression positively correlated with increasing body weight in humans. These findings reveal a surprising relationship between aTregs and adipocytes in promoting insulin resistance during excessive caloric intake, placing Blimp-1-regulated IL-10 expression by aTregs at a critical juncture in the development of obesity and its associated comorbidities in mice and humans.

**SUMMARY:** Here we show that ablation of Blimp-1 in adipose tissue resident Tregs (aTregs) leads to decreased IL-10 production, resulting in increased Ucp-1 expression and beiging by adipocytes and protection from diet-induced obesity and insulin resistance.

## INTRODUCTION

Rising obesity levels worldwide have encouraged investigators to think more broadly when determining the factors that can regulate visceral adipose tissue (VAT) adipocytes and the development of chronic low-grade inflammation associated with obesity(*1*). It is now clear that the innate and adaptive immune systems play critical roles in protection from obesity and associated inflammation(*2*). Visceral adipose tissue resident Tregs (aTregs) are a substantial population of Foxp3^+^ CD4^+^ T cells that express the signature transcription factor PPAR*γ*, produce the cytokine interleukin-10 (IL-10) and appear to be critical for the suppression of obesity-associated inflammation(*3–5*). In humans, as in mice, increased body mass due to diet-induced obesity (DIO) corresponds with decreased frequency and number of aTregs(*4*). Although extensive investigation has revealed key genes and functions of aTregs (*3, 4, 6, 7*), it is unclear how aTregs communicate with adipocytes, the predominant cell type in the VAT.

White adipose tissue (WAT) is primarily composed of adipocytes, preadipocytes, stromal cells and immune cells(*8, 9*). The main function of adipocytes is to store lipids which can be converted to fatty acids (FAs) for use by liver, heart and skeletal muscle cells, among others, as required. However, adipocytes can also secrete adipokines, such as leptin and adiponectin, to modulate metabolism and VAT-resident immune cells(*10*). Brown adipose tissue (BAT), in contrast, is required for thermogenesis and heat generation upon cold exposure and loss of BAT is associated with increased weight gain(*11, 12*). Recent data have shown that certain immune cells including innate lymphoid cells (ILCs) and eosinophils can produce IL-4, which acts on adipocytes in WAT to increase beiging and thermogenesis in these cells(*13, 14*). Immune-modulated IL-4 expression results in upregulation of Uncoupling Protein 1 (Ucp-1) which positively regulates thermogenesis and energy expenditure in adipocytes(*13, 14*). Whether aTregs can also participate in the regulation of beiging in adipocytes is the focus of this study.

B lymphocyte-induced maturation protein-1 (Blimp-1) is a zinc finger motif containing transcriptional regulator encoded by the gene positive regulatory domain 1 (*prdm1*)(*15, 16*). Blimp-1 is known to be involved in the development, polarization, and maintenance of various immune cell types including development of B cells and differentiation of effector CD8^+^ T cells(*16–18*). In mucosal resident effector Tregs, deletion of Blimp-1 results in a significant loss of IL-10(*19*). Interestingly, in adipose tissue Tregs, Blimp-1 is constitutively high(*7, 19*), and correlates with ST2 expression and IL-10 production(*7*). Given the positive association between Blimp-1 and IL-10 in Tregs, we decided to explore whether Blimp-1^+^ IL-10-producing aTregs could modulate adipocyte function under normal and obese conditions. IL-10 expression often correlates with reduced tissue inflammation(*20*), leading us to postulate that loss of Blimp-1 driven IL-10 expression in aTregs would result in increased VAT inflammation, DIO and insulin resistance.

Surprisingly, we show here that although Treg-specific Blimp-1 deficiency resulted in reduced IL-10 production by aTregs, this did not result in the expected increased inflammation and insulin resistance that we expected. Instead, we observed increased expression of Ucp-1 by adipocytes, and increased weight loss, whole-body fat oxidation and insulin sensitivity. Moreover, increased IL-10 expression correlated with increasing BMI in humans, placing Blimp-1 dependent IL-10 production by aTregs at the center of crosstalk between aTregs and adipocytes.

## RESULTS

### Blimp-1 is expressed in a subset of aTregs

Previous data have shown that Blimp-1 expression in aTregs is high, relative to splenic Tregs, and suggested a correlation between Blimp-1 and the aTreg marker ST2, the IL-33 receptor(*7*). We first confirmed that Blimp-1 was expressed by the majority of aTregs, as previously reported(*7*). To this end, we crossed Foxp3-RFP mice to Blimp-1-YFP BAC-transgenic reporter mice. In contrast to Tregs in the spleen, approximately 50% of all aTregs expressed Blimp-1 (Fig. 1A). As Blimp-1 has been described as a critical transcriptional regulator of effector Tregs in peripheral tissues(*21*), we next assessed whether Blimp-1 expression in aTregs correlated with an ‘effector-like’ phenotype in aTregs. Our analysis showed that Blimp-1^+^ aTregs expressed increased levels of ST2 and KLRG1, indicating that Blimp-1 expression marks terminal differentiated aTregs (Fig. 1B). Blimp-1 has been previously reported to repress CD25 expression on CD8^+^ effector T cells(*22*). Similarly, we noted that CD25 expression was reduced in Blimp-1^+^ aTregs (Fig. 1C). Together, these data confirm that Blimp-1 expression marks aTregs that exhibit an effector-like Treg phenotype relative to Blimp-1^-^ aTregs, and that Blimp-1^+^ Tregs were enriched in the VAT relative to Tregs in the spleen.

**Figure 1.**
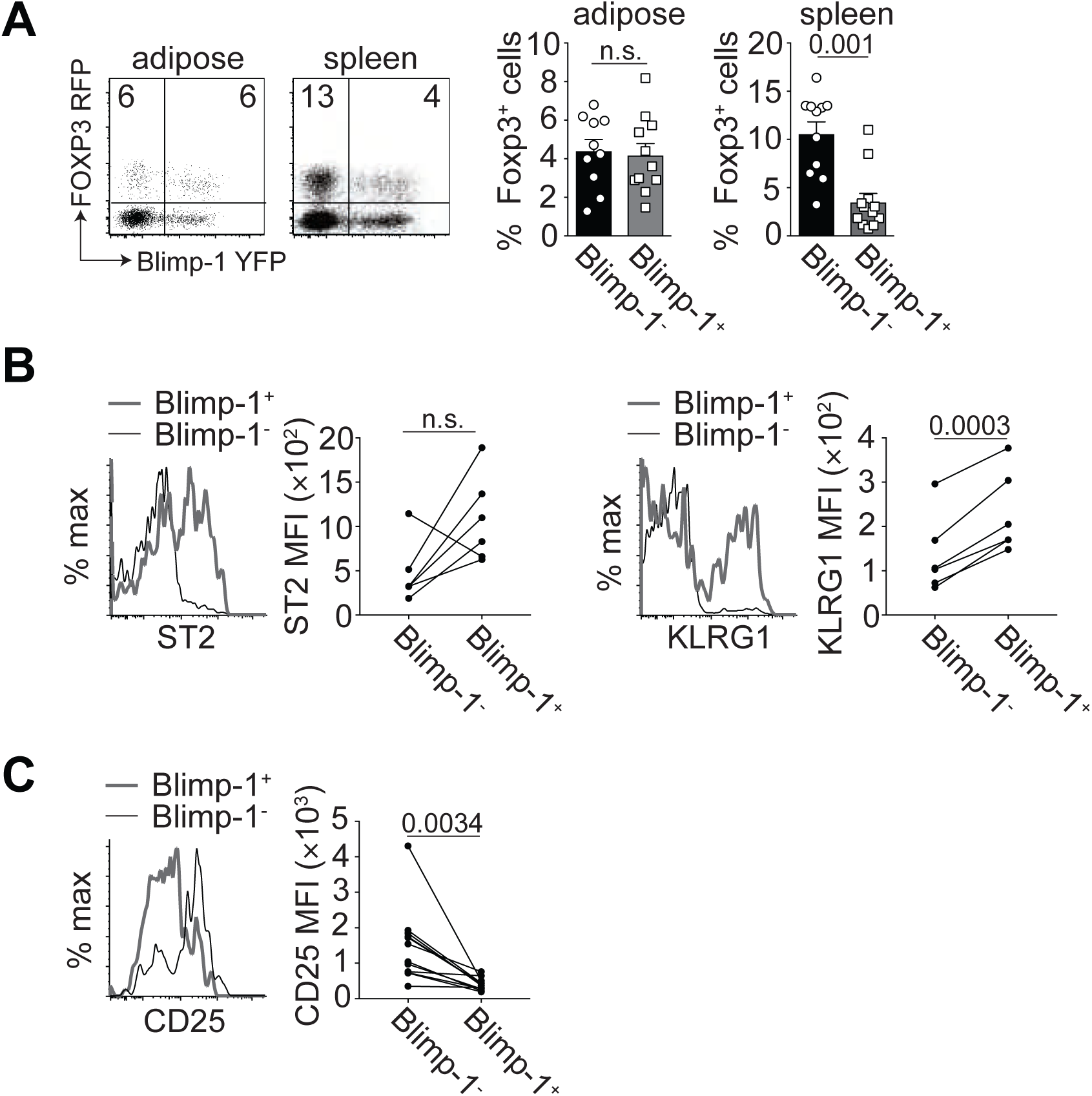
Blimp-1 expression in Tregs from the VAT and spleen. Cells were isolated from the spleen or visceral adipose tissue (VAT) of 15-week-old male mice expressing Blimp-1-YFP and Foxp3-RFP. (A) Flow cytometry plots and bar graphs indicating the frequency of CD4^+^ gated Blimp-1^+^ and Blimp-1^-^ Foxp3^+^ Tregs from the indicated tissues. Each dot represents one animal. (B) Histograms and graphs indicating ST2 and KLRG1 expression on gated CD4^+^ Foxp3^+^ aTregs that were either Blimp-1^+^ (grey line) or Blimp-1^-^ (black line). (C) Histograms and graphs indicating CD25 expression on gated CD4^+^ Foxp3^+^ aTregs that were either Blimp-1^+^ (grey line) or Blimp-1^-^ (black line). Data are presented as means ± s.e.m. for *n* = 11 mice per group (A) or *n* = 6 mice per group (B and C), pooled from four independent experiments. A paired Student’s *t*-test was performed to determine significance and the *P* values are indicated on the graphs (n.s. = not significant).

### Phenotype of Blimp-1 deficient aTregs in diet-induced obesity

To assess the function of Blimp-1 expression in aTregs, we crossed transgenic mice expressing Cre recombinase and YFP under the control of Foxp3 regulatory elements to mice carrying conditional (loxp-flanked) *prdm1* alleles to generate Blimp-1^f/f^ Foxp3-Cre^+^ mice. We then probed how the aTreg population was affected by loss of Blimp-1 expression, relative to Tregs in the spleen. Using 26-28-week-old Blimp-1^f/f^ Foxp3-Cre^+^ male mice on standard fat diet (SFD), we observed that loss of Blimp-1 expression resulted in a significant decrease in the frequency of Foxp3^+^ aTregs, while Tregs in the spleen were unaffected (fig. S1, A). Furthermore, Blimp-1 aTreg deficiency resulted in decreased expression of ST2, CCR2, GITR and KLRG1 in aTregs, all markers of aTregs (fig. S1, B).

We next placed Blimp-1^f/f^ Foxp3-Cre^+^ male mice on 60% HFD for 18-20 weeks. We determined that the overall frequency and number of aTregs was unperturbed relative to WT mice, although the frequency of Tregs in the spleen was significantly increased in the Blimp-1^f/f^ Foxp3-Cre^+^ animals (Fig. 2A). As expected in mice on HFD, the frequency and number of aTregs was reduced relative to Tregs in the spleen, consistent with previously published data (Fig. 2A)(*4*). Phenotypically, loss of Blimp-1 expression in aTregs resulted in decreased ST2, CCR2, GITR and PD1 expression (Fig. 2B), while KLRG1 expression was not significantly reduced (data not shown). Consistent with these data, loss of Blimp-1 expression in all T cells using a CD4-Cre line similarly showed reduced expression of ST2, KLRG1 and CCR2, while the frequency and number of aTregs was unchanged (fig. S2, A and B). Lastly, we tested expression of known aTreg-associated transcription factors in the absence of Blimp-1. Although we did not detect a change in GATA3 expression in the absence of Blimp-1 (data not shown), we observed that the aTreg signature gene, PPAR*γ*, was significantly down regulated, resulting in fewer PPAR*γ*^+^ aTregs and lower PPAR*γ* expression on a per cell basis (Fig. 2C). Overall, our data were consistent with a role for Blimp-1 in positive regulation of markers associated with aTreg differentiation.

**Figure 2.**
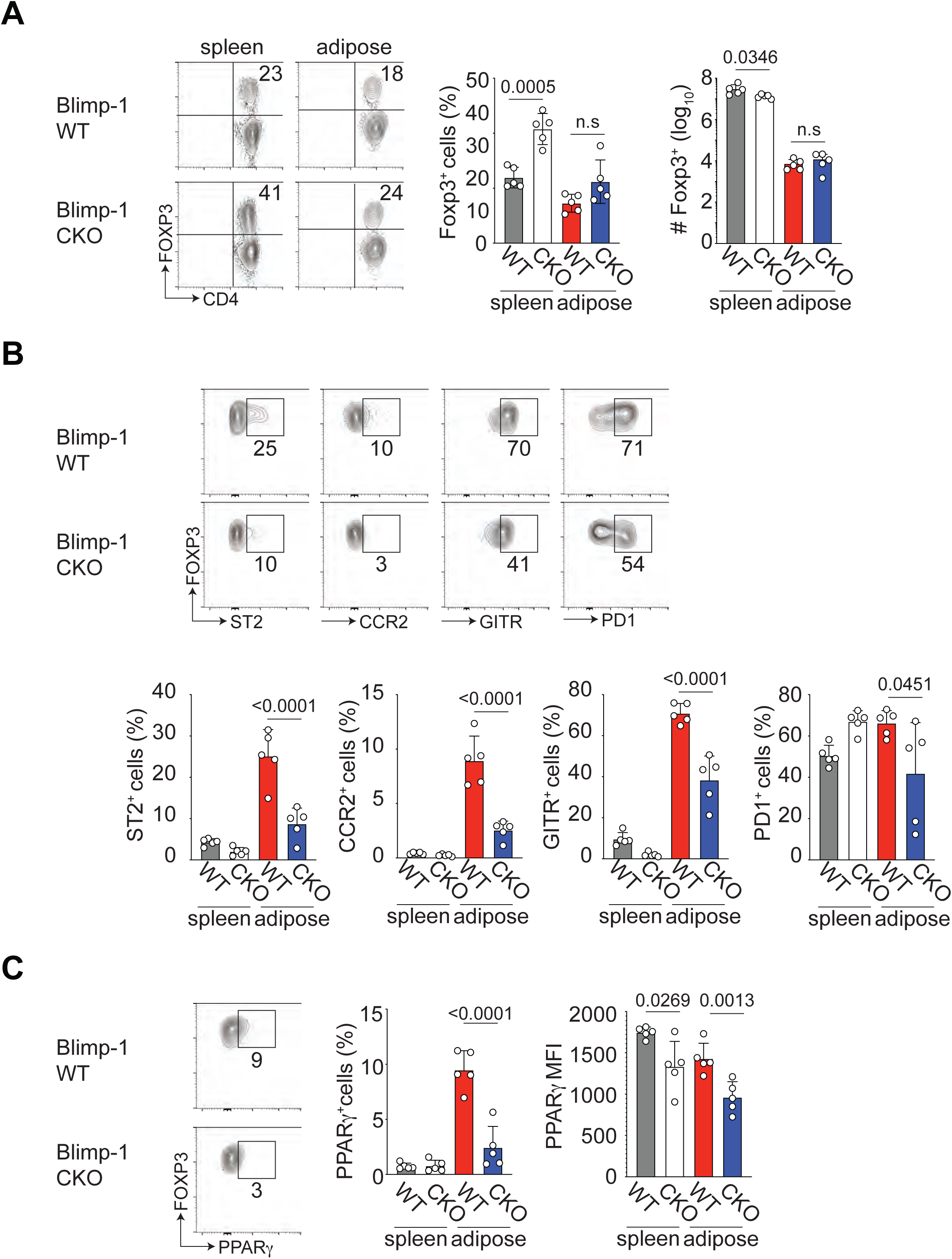
Characterization of Blimp-1 deficient Tregs from mice on high fat diet (HFD). Male Foxp3-YFP-Cre^+^ (WT) and Blimp-1^f/f^ mice crossed to Foxp3-YFP-Cre^+^ (conditional knockout, CKO) were placed on 60% high fat diet (HFD) for 18 weeks prior to metabolic analysis. (A) Flow cytometry plots and bar graphs showing the frequency and number of CD4^+^ Foxp3^+^ cells in the indicated tissue from WT and CKO mice. (B) Flow cytometry plots and bar graphs showing expression of ST2, CCR2, GITR and PD1 on gated CD4^+^ Foxp3^+^ cells in the indicated tissue from WT and CKO mice. (C) Flow cytometry plots and bar graphs showing expression of PPAR*γ* on gated CD4^+^ Foxp3^+^ cells as a percentage and median fluorescent intensity (MFI) in the indicated tissue from WT and CKO mice. Each dot represents one animal. Data are presented as means ± s.e.m. for *n* = 5 mice per group, pooled from at least 2 independent experiments. A paired Student’s *t*-test was performed to determine significance and the *P* values are indicated on the graphs (n.s. = not significant).

### Blimp-1 expression in aTregs promotes insulin resistance and obesity

Given that the aTreg phenotype was perturbed in the absence of Blimp-1 under SFD and HFD conditions, we next wanted to assess how Blimp-1^+^ aTregs affected insulin sensitivity and glucose tolerance. Analysis of the total body weight, fasting plasma insulin and blood glucose levels in 26-28-week-old SFD-fed male mice did not reveal any significant difference (fig. S3, A-C). However, we noted a surprising trend towards decreased body weight, and fasting insulin and glucose levels in the Blimp-1^f/f^ Foxp3-Cre^+^ mice relative to the WT controls (fig. S3, A-C). Similarly, when we performed a glucose tolerance test (GTT) on WT and Blimp-1^f/f^ Foxp3-Cre^+^ mice, we observed decreased plasma insulin and blood glucose levels in Blimp-1^f/f^ Foxp3-Cre^+^ mice compared with WT mice over time (fig. S3, D and E).

We next placed 8-week old male mice on 60% fat diet (HFD) for 18-20 weeks prior to our analysis. We measured the overall body weight of the Blimp-1^f/f^ Foxp3-Cre^+^ animals relative to Foxp3-Cre^+^ only (WT) mice after HFD. Unexpectedly, we observed a consistent decrease in the overall body weight of Blimp-1^f/f^ Foxp3-Cre^+^ mice (Fig. 3A). Furthermore, fasting insulin and glucose levels in the Blimp-1^f/f^ Foxp3-Cre^+^ mice were significantly reduced (Fig. 3, B and C), such that the HOMA-IR, a measure of insulin resistance, was substantially reduced in the Blimp-1^f/f^ Foxp3-Cre^+^ mice (Fig. 3D). There data suggested that loss of Blimp-1 expression in Tregs was protective against diet-induced insulin resistance. Consistent with these data, plasma insulin and blood glucose levels were reduced in GTTs in Blimp-1^f/f^ Foxp3-Cre^+^ mice relative to WT controls, demonstrating improved glucose tolerance (Fig. 3, E and F). Moreover, weight gain over time on HFD was consistently reduced in the Blimp-1^f/f^ Foxp3-Cre^+^ mice (Fig. 3G and fig. S4). Analysis of the colon length in the HFD-fed Blimp-1^f/f^ Foxp3-Cre^+^ mice did not reveal any gross abnormalities (Fig. 3H), and thus we concluded that the reduced weight gain and improved insulin sensitivity we observed in the HFD-fed Blimp-1^f/f^ Foxp3-Cre^+^ mice was not due to colitis or nutrient malabsorption in the gut. In summary, these data showed that loss of Blimp-1 expression in Tregs can substantially reduce weight gain and fasting blood glucose and plasma insulin levels, as well as enhance insulin sensitivity in HFD-fed mice.

**Figure 3.**
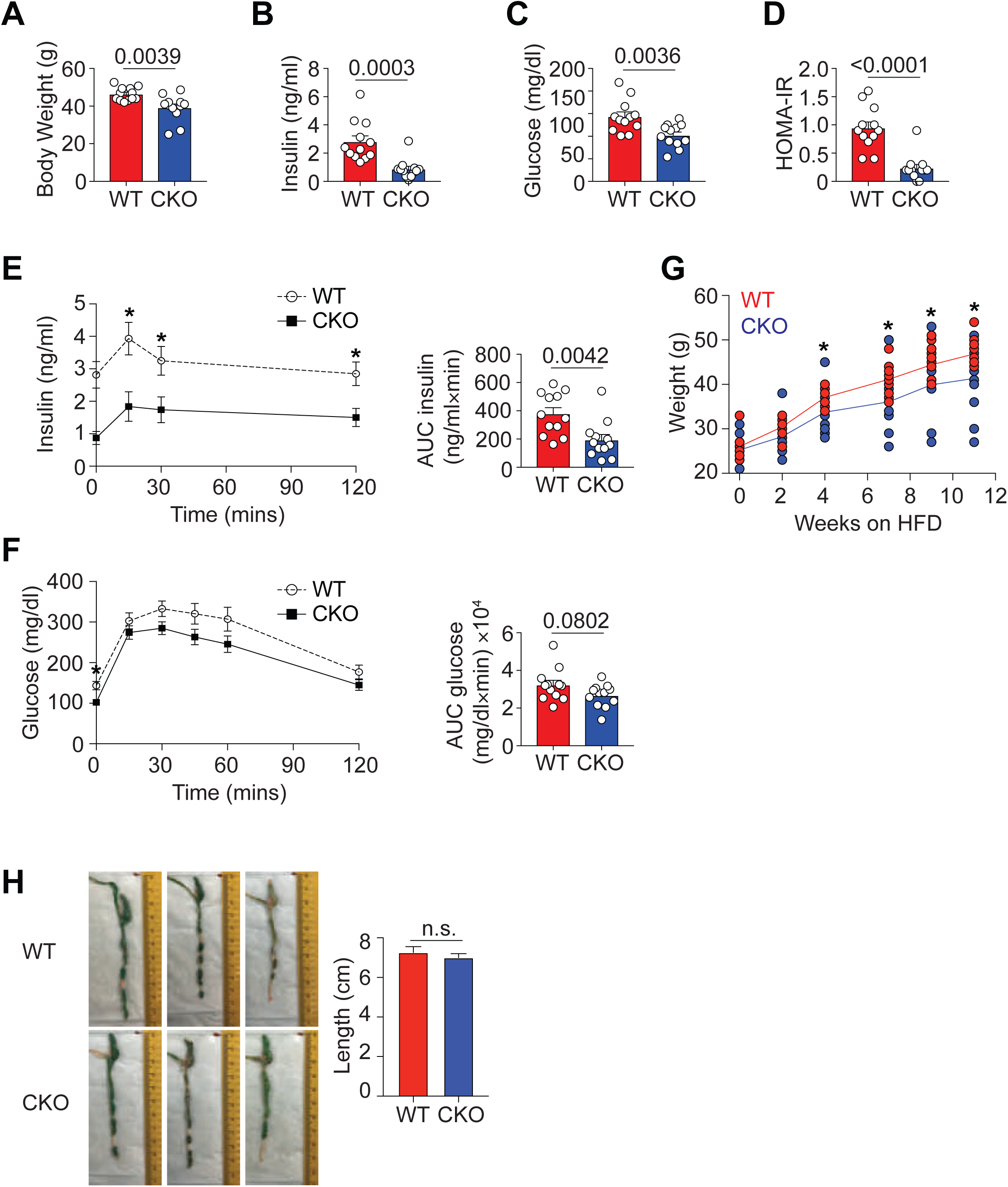
Loss of Blimp-1 expression in Tregs decreases weight and increases insulin sensitivity. Male Foxp3-YFP-Cre^+^ (WT) and Blimp-1^f/f^ mice crossed to Foxp3-YFP-Cre^+^ (conditional knockout, CKO) were placed on 60% high fat diet (HFD) for 18 weeks prior to metabolic analysis. (A) Bar graphs indicating body weight of 26-week old HFD-fed WT and CKO mice. (B) Bar graph showing fasting plasma insulin levels in WT and CKO mice. (C) Bar graph showing fasting glucose blood levels in WT and CKO mice. (D) Bar showing the homeostatic model assessment of insulin resistance (HOMA-IR) in WT and CKO mice. (E) Graph indicating plasma insulin levels in mice over time after i.p. glucose injection. Bar graph indicates the area under the curve (AUC) for both groups. (F) An i.p. glucose tolerance test (GTT) was performed on WT and CKO mice in (E). Bar graph indicates the area under the curve (AUC) for both groups. (G) Graph indicating total body weight per mouse per group over time on HFD. (H) Photographs and quantification of the colon length in WT and CKO mice after 18-20 weeks on HFD. For all experiments each dot represents one mouse and an unpaired Student’s *t*-test was performed to determine significance (A-D). Comparisons at each time point were made against WT control mice by ANOVA where *p<0.05 (E, G). Data are presented as means ± s.e.m., for *n* = 12 mice per group, pooled from at least 2 independent experiments.

### Blimp-1 deficiency in Tregs decreases fat mass and food intake

We next determined the influence of Treg-specific Blimp-1 expression on energy homeostasis in SFD- and HFD-fed mice. Male mice at 26 weeks of age were individually housed in metabolic chambers for 48 hours. MRI analysis of body composition confirmed that Blimp-1^f/f^ Foxp3-Cre^+^ mice had lower total body weight that was due primarily to reduced fat mass relative to the WT controls. As a result, lean mass as a percentage of total body weight was increased and fat mass as a percentage of body weight was reduced in Blimp-1^f/f^ Foxp3-Cre^+^ mice, irrespective of whether the mice were on SFD or HFD (fig. S3, F-H, Fig. 4, A-C). Remarkably, in SFD-fed mice, food consumption, expressed per kg lean mass, was increased in the Blimp-1^f/f^ Foxp3-Cre^+^ mice (fig. S3, I) but our data showed that this increased food consumption did not result in increased weight gain in the SFD group (fig. S3, F-H). Moreover, the respiratory exchange ratio (RER) was increased in the Blimp-1^f/f^ Foxp3-Cre^+^ mice, relative to the control group, indicating these animals increased whole body glucose oxidation relative to fat oxidation (fig. S3, J).

**Figure 4.**
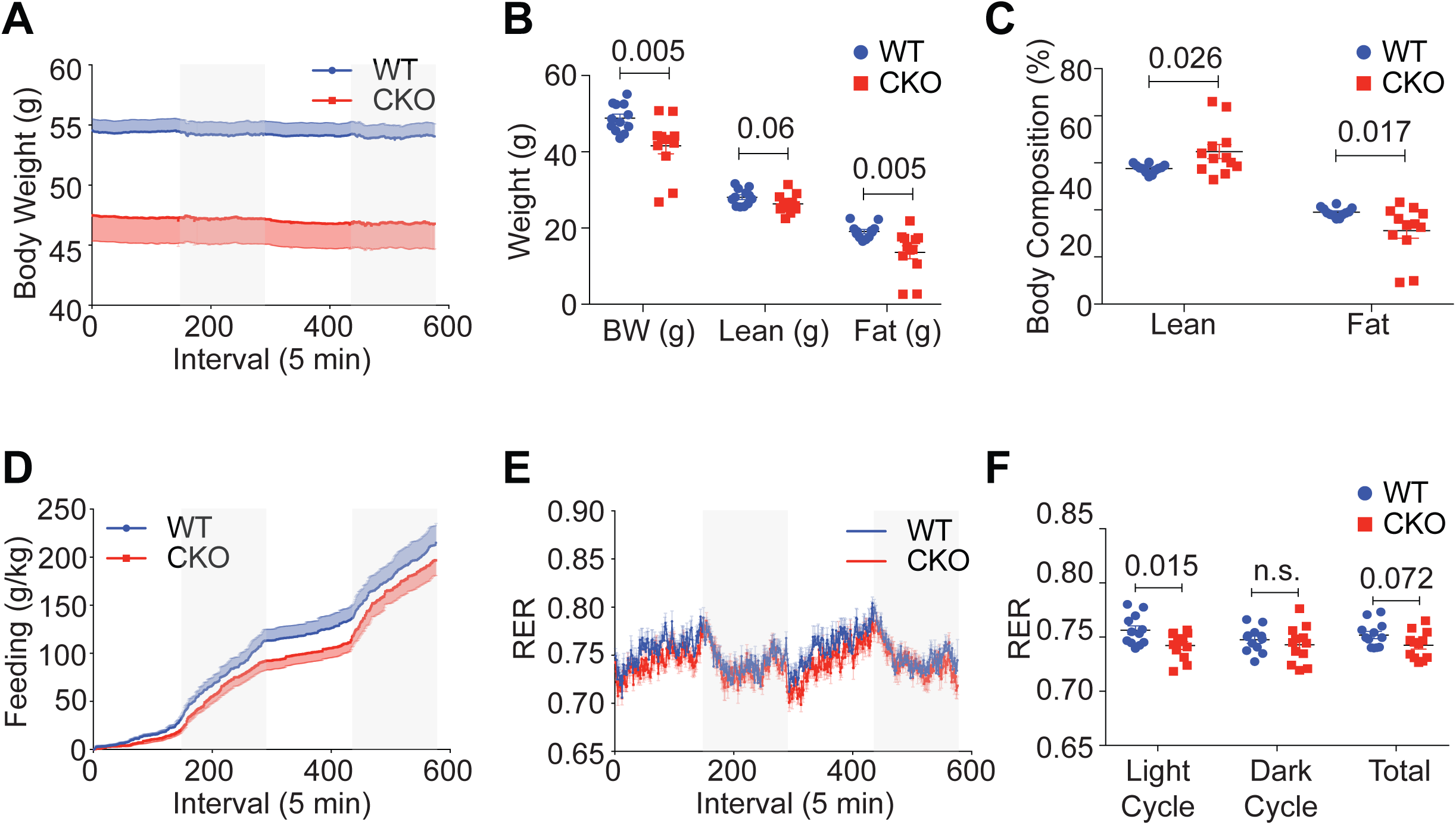
Calorimetry and body composition changes with loss of Blimp-1 expression in Tregs. Male Foxp3-YFP-Cre^+^ (WT) and Blimp-1^f/f^ mice crossed to Foxp3-YFP-Cre^+^ (conditional knockout, CKO) were placed on 60% high fat diet (HFD) for 18 weeks prior to metabolic analysis. (A) Graph showing body weight as measured by Promethion Multiplexed Metabolic Cage System for 48-hr total duration in WT and CKO mice. (B) Body weight (BW), lean and fat mass in grams for WT and CKO mice. (C) Percent lean and fat mass expressed per gram body weight in WT and CKO mice. (D) Food intake in g/kg lean mass for WT and CKO mice. (E) Respiratory exchange ratio (RER) over time in WT and CKO mice. (F) Respiratory exchange ratio (RER) in light, dark and total in WT and CKO mice. Data are presented as means ± s.e.m., for *n* = 12 mice per group, pooled from at least 2 independent experiments. Each dot represents one mouse and an unpaired Student’s *t*-test was performed to determine significance.

In HFD-fed mice, food consumption expressed per kg lean mass was modestly reduced in the Blimp-1^f/f^ Foxp3-Cre^+^ mice (Fig. 4D) and the RER during the light cycle was reduced in the absence of Blimp-1 Treg expression (Fig. 4E, F). The reduced RER in the Blimp-1^f/f^ Foxp3-Cre^+^ mice during the light cycle indicated increased whole-body fat oxidation, which may have contributed to the reduced fat mass in the Blimp-1-deficient mice on HFD. Taken together, our data show that SFD-fed Blimp-1^f/f^ Foxp3-Cre^+^ mice weigh less and consume more calories while HFD-fed Blimp-1^f/f^ Foxp3-Cre^+^ mice weigh less and consume fewer calories, resulting in SFD- and HFD-fed mice that are protected from DIO.

### Global immune landscape in the VAT with loss of Blimp-1 in Tregs

Given our contradictory data showing that loss of Blimp-1 expression in aTregs promoted protection from DIO but also resulted in reduced differentiation of aTregs, we postulated that loss of ST2 expression on aTregs enabled other ST2-positive immunoregulatory cells in the VAT, such as ILC2s and M2 macrophages, to respond to the increased IL-33, and thus promote insulin sensitivity and weight loss. However, we did not observe a difference in the frequency of total CD45^+^ cells from the stromal vascular fraction (SVF) or indeed a difference in the frequency of aTreg^-^ ST2^+^ CD45^+^ cells in the Blimp-1^f/f^ Foxp3-Cre^+^ mice relative to WT mice (Fig. 5A), leading us to conclude that the increased insulin sensitivity and weight loss we observed in Blimp-1^f/f^ Foxp3-Cre^+^ mice was not due to expansion of other ST2^+^ immunoregulatory cells in the VAT.

**Figure 5.**
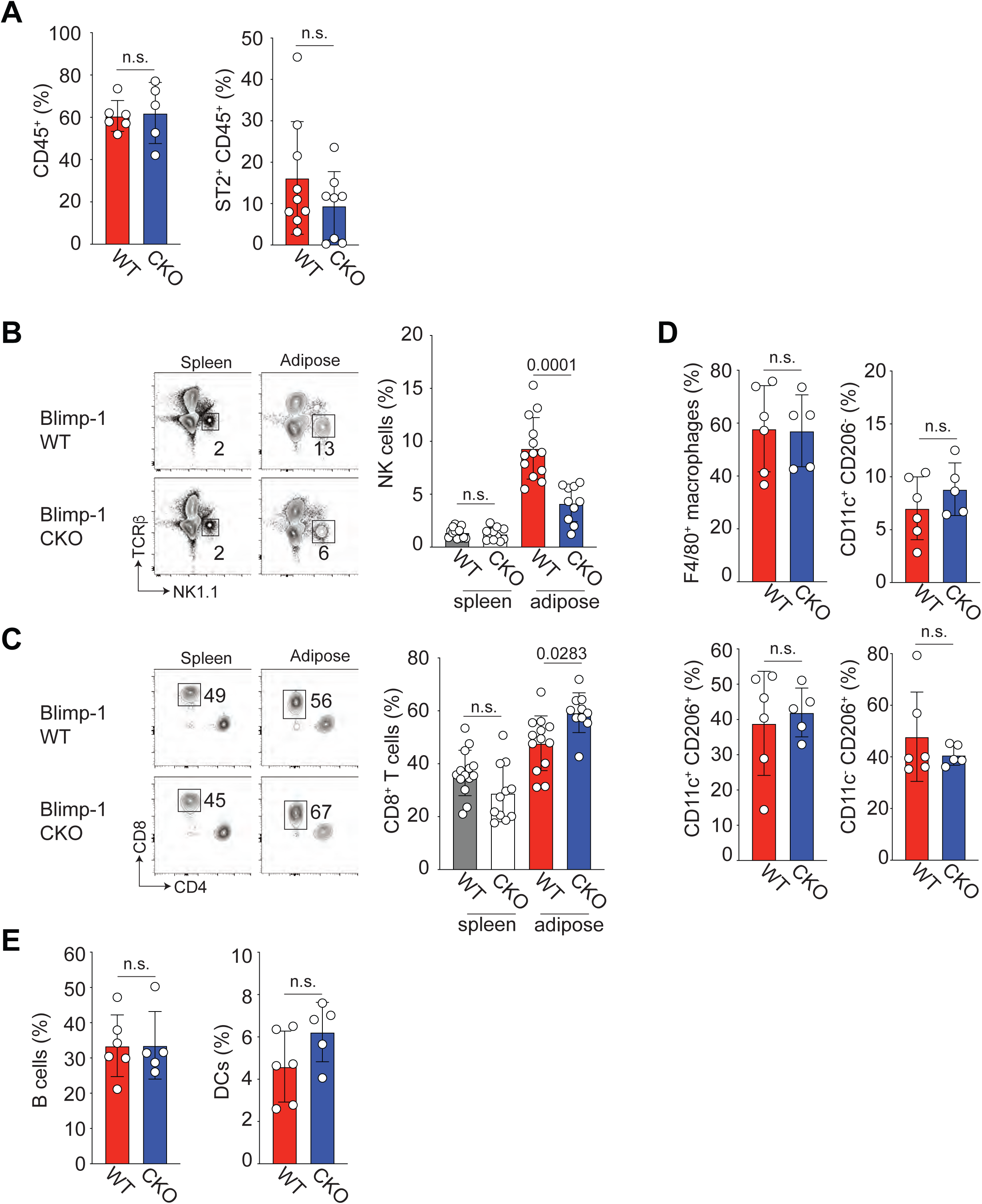
Global immune-phenotyping of VAT in Blimp-1 aTreg-deficient animals. Male Foxp3-YFP-Cre^+^ (WT) and Blimp-1^f/f^ mice crossed to Foxp3-YFP-Cre^+^ (conditional knockout, CKO) were placed on 60% high fat diet (HFD) for 18 weeks prior to global immune-phenotyping analysis. (A) Bar graphs indicating the frequency of total CD45^+^ cells (left) and aTreg^-^ ST2^+^ CD45^+^ cells (right) isolated from the stromal vascular fraction (SVF) of WT or CKO mice on HFD. (B) Flow cytometry plots and bar graphs showing the frequency of NK1.1^+^ cells from gated CD45^+^ live^+^ cells isolated from the spleen or VAT of WT and CKO mice. (C) Flow cytometry plots and bar graphs showing the frequency of CD8^+^ cells from gated CD45^+^ live^+^ cells isolated from the spleen or VAT of WT and CKO mice. (D) Bar graphs showing the frequency of the indicated macrophage populations isolated from the VAT of WT or CKO mice. (E) Bar graphs showing the frequency of B cells and dendritic cells (DCs) isolated from the VAT of WT or CKO mice. Data are presented as means ± s.e.m., for *n* = 6-14 mice per group, pooled from at least 2 independent experiments. Each dot represents one mouse and an unpaired Student’s *t*-test was performed to determine significance.

Because loss of Blimp-1 expression in aTregs in HFD-fed mice had perturbed aTreg differentiation and resulted in protection of these animals from DIO, we next tested if any other immune cell populations had also been affected by loss of Blimp-1 expression in the aTregs. We noted that the frequency of natural killer (NK) cells was significantly decreased in the VAT, while the frequency of CD8^+^ T cells was slightly increased (Fig. 5, B and C). However, the frequencies of macrophage subsets, B cells and dendritic cells (DCs) were all unaffected by loss of Blimp-1 expression in aTregs (Fig. 5, D and E). Overall, we show here that most immune cell populations are unaffected by loss of Blimp-1 expression in aTregs. VAT-resident NK cells have previously been associated with increased insulin resistance, indicating that a reduction in NK cell frequency(*14, 23, 24*) likely contributed to the reduced insulin resistance and weight loss we observed in the Blimp-1^f/f^ Foxp3-Cre^+^ mice. Although CD8^+^ T cells are inflammatory and are recruited to the VAT of HFD-fed mice(*25*), we concluded from our data that this increase in VAT CD8^+^ T cells was too slight to modulate the significant protection from weight gain and insulin resistance we observed in the Blimp-1^f/f^ Foxp3-Cre^+^ mice.

### Blimp-1 regulated IL-10 production by aTregs represses Ucp-1 in adipocytes

Given that the distinctive weight loss and insulin sensitivity we observed in Blimp-1^f/f^ Foxp3-Cre^+^ mice on SFD and HFD could not be adequately explained by changes in the other immune cell populations in the VAT, we next determined if distinct gene expression changes by the Blimp-1-deficient aTregs could be responsible for these phenomena. We performed NextGen RNA-sequencing on WT Blimp-1^f/f^ Foxp3-Cre^+^ aTregs from SFD-fed mice and compared gene expression changes in these two populations, relative to Tregs sorted from the spleen. We noted that a number of aTreg-associated genes were downregulated in the Blimp-1-deficient aTregs, including Il-10, Il1rl1 (ST2), Klrg1 and Ppar*γ* (Fig. 6A). As direct positive regulation of IL-10 production by Blimp-1 in splenic Tregs was previously described(*19, 21*), this prompted us to test whether IL-10 protein expression was reduced in aTregs in the adipose tissue. Isolation of aTregs from the stromal vascular fraction (SVF) of the VAT in WT and Blimp-1^f/f^ Foxp3-Cre^+^ mice revealed that IL-10 expression was significantly reduced in Blimp-1 deficient aTregs in mice on SFD and HFD (fig. S5, Fig. 6B). We also tested expression of TNF*α* and IFN*γ* from Blimp-1 deficient aTregs and noted that these inflammatory cytokines were increased in aTregs in the absence of Blimp-1 (fig. S6, A and B). Although inflammatory cytokines typically help to recruit inflammatory immune cells such as M1 macrophages to the adipose tissue, we determined there were no differences in these cell populations (Fig. 5, D and E), and thus that the effect of inflammatory cytokine production by Blimp-1-deficient aTregs was negligible. Additionally, it was previously shown that TNF*α* can help to drive IL-33 expression by VAT stromal cells(*26*), and therefore increased TNF*α* expression by Blimp-1-deficient aTregs could help with this process.

**Figure 6.**
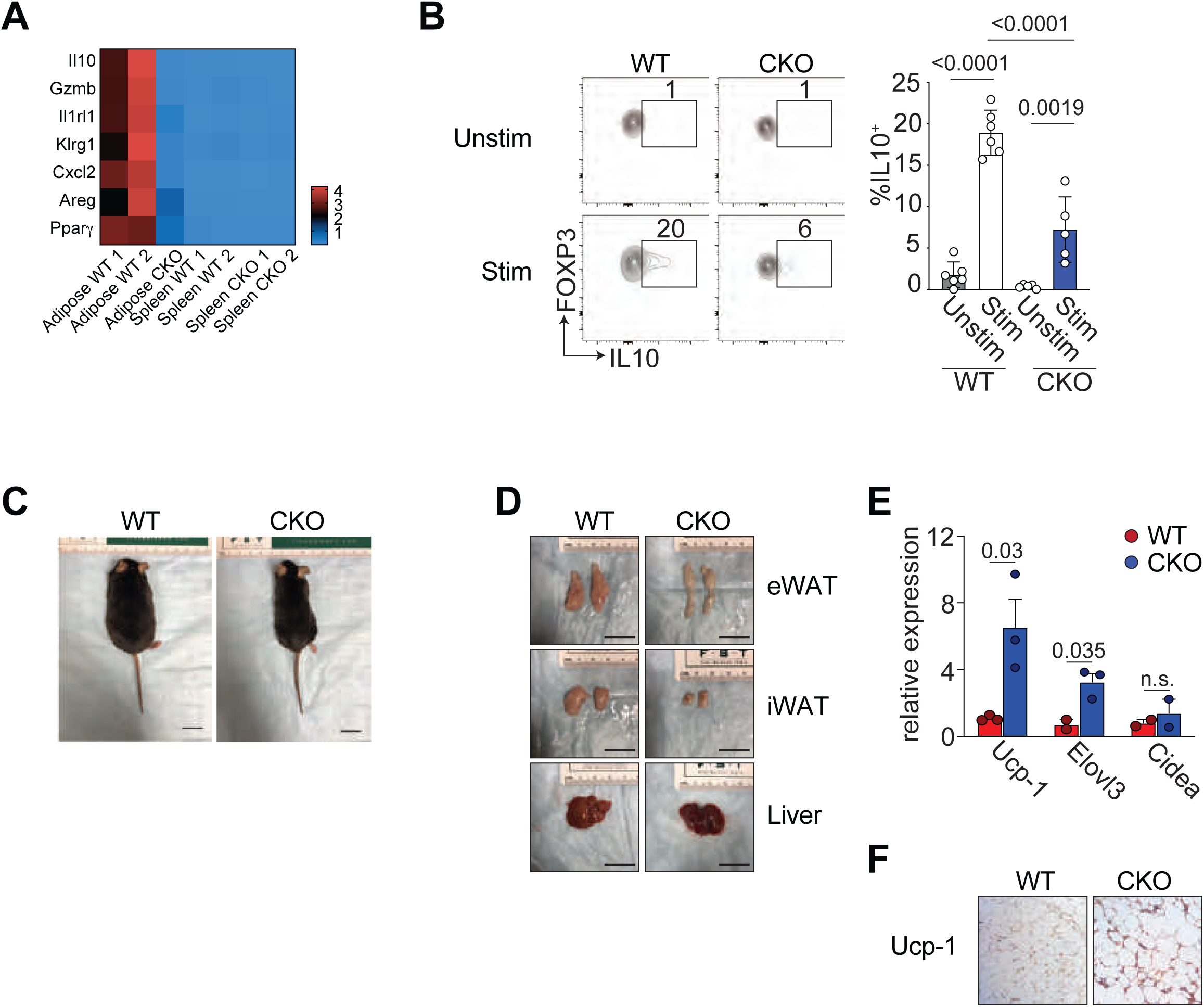
Loss of IL-10 expression by Blimp-1-deficient aTregs increases Ucp-1 expression in visceral adipocytes. (A) Heatmap representing select aTreg-associated genes that were differentially expressed in WT and CKO aTregs. (B) Flow cytometry plots and bar graphs showing expression of IL-10 by gated CD4^+^ Foxp3^+^ cells that were unstimulated (unstim) or stimulated (stim) for 5 hours with PMA and ionomycin from the VAT of WT and CKO mice. (C) Gross appearance of 26-28-week old HFD-fed WT and CKO mice. (D) Gross appearance of the epididymal WAT (eWAT), inguinal WAT (iWAT) and livers from 26-28-week old HFD-fed WT and CKO mice. (E) Relative mRNA expression of Ucp1, Elovl3 and Cidea in WT and CKO VAT isolated from 26-28-week old HFD-fed WT and CKO mice. Data were normalized to HPRT. (F) Immunohistochemical staining for Ucp-1 in adipocytes from the epidydimal white adipose tissue (eWAT). Data are presented as means ± s.e.m., for *n* = 5-6 mice per group, pooled from at least 2 independent experiments (B) or representative from at least 2 experiments with 5 mice per group (C) and (D). Each dot represents one mouse in (E) and an unpaired Student’s *t*-test was performed to determine significance (n.s. = not significant). Each dot in (B) represents one mouse and one-way ANOVA was performed to determine significance. The scale bar for (C) and (D) is equal to 2 cm.

IL-10 signaling was recently shown to be a critical repressor of adipocyte beiging and thermogenesis(*11*). We thus reasoned that reduced IL-10 production from aTregs in the absence of Blimp-1 would result in decreased fat mass and increased expression of Ucp-1, a protein typically expressed in brown adipocytes and responsible for energy expenditure and thermogenesis(*11*), in adipocytes from the WAT. HFD-fed Blimp-1^f/f^ Foxp3-Cre^+^ mice were grossly leaner than WT controls (Fig. 6C), indicating potentially increased beiging of WAT in the Blimp-1-deficient animals. Furthermore, visual examination of the epididymal WAT (eWAT) and the inguinal WAT (iWAT) fat pads, revealed these fat pads to be smaller than those from WT mice, although we could detect no overt differences between the livers from the two groups (Fig. 6D).

As loss of IL-10 signaling in white adipocytes results in increased expression of Ucp-1 expression, we next assessed expression of Ucp-1 and other thermogenic genes by adipocytes in the Blimp-1^f/f^ Foxp3-Cre^+^ mice. We detected significantly increased mRNA expression of Ucp-1 and Elovl3 in adipocytes isolated from Blimp-1^f/f^ Foxp3-Cre^+^ mice relative to WT controls (Fig. 6E). Additionally, we observed increased Ucp-1 protein expression in the eWAT of Blimp-1^f/f^ Foxp3-Cre^+^ mice relative to their WT counterparts (Fig. 6F), indicating that loss of IL-10 production from the Blimp-1-deficient aTregs resulted in increased Ucp-1 expression in the adipocytes of these animals, and that this increased energy expenditure by the white adipocytes resulted in increased weight loss and insulin sensitivity during DIO.

### IL-10 production in aTregs in humans positively correlates with increasing BMI

To determine whether IL-10 expression correlated with increasing body mass index (BMI) in humans, we performed metadata analysis on publicly available data from the Metabolic Syndrome in Men (METSIM) project. In this study, abdominal subcutaneous adipose tissue from 434 male subjects was analyzed by RNA-seq (GEO Accession Number: GSE135134). Correlating BMI with *FOXP3* transcripts expressed in T cells, we observed as previously reported(*4*), that *FOXP3* transcripts negatively correlated with increasing BMI (Fig. 7A). Similarly, as recently published, we observed that expression of IL-10R transcripts increased with increasing BMI (Fig. 7B)(*11*). Although expression of Blimp-1 did not positively corelate with increasing BMI (data not shown), we noted that as BMI increased, IL-10 transcripts in the adipose tissue increased, even as expression of Foxp3 decreased (Fig. 7, C and D). Thus, these data suggest that, in humans, increasing IL-10 expression positively correlates with increasing BMI and suggest that aTreg-produced IL-10 could be critical for repressing white adipocyte thermogenesis and increasing insulin resistance in humans in a manner similar to our observations in mice.

**Figure 7.**
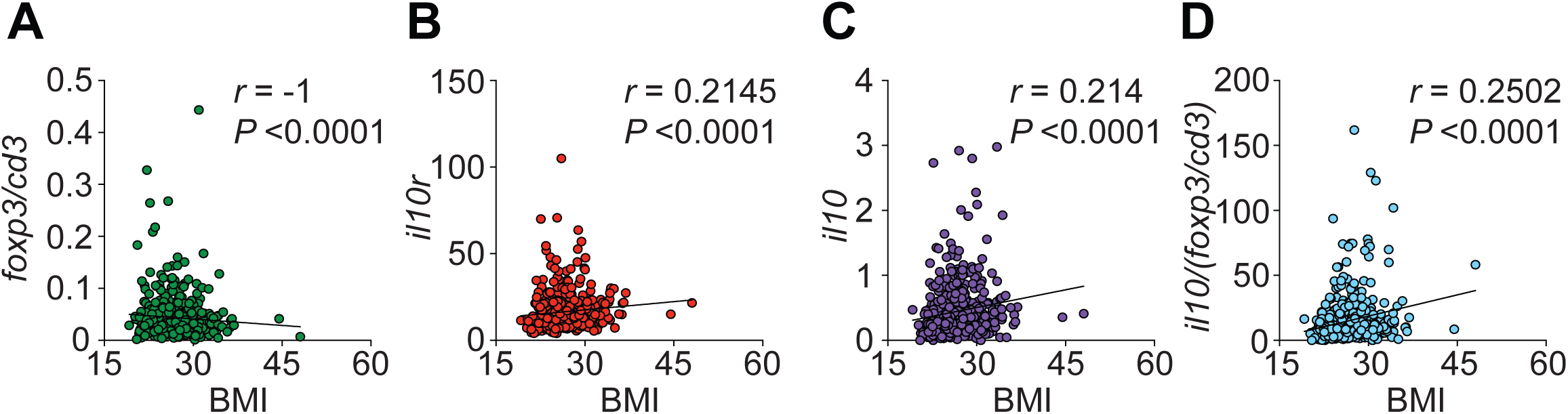
IL-10 expression in adipose tissue increases with BMI in humans. Using the METSIM project, transcripts per million [TPM] mRNA was correlated by linear regression with increasing BMI using Spearman’s correlation coefficient. (A) Correlation trait plots of Foxp3/CD3 mRNA ratio and increasing BMI. (B) Correlation trait plots of IL-10r mRNA and increasing BMI. (C) Correlation trait plots of IL-10 mRNA and increasing BMI. (D) Correlation trait plots of IL-10/(Foxp3/CD3) mRNA ratio and increasing BMI. *n* = 434 human males, GEO Accession Number: GSE135134.

## DISCUSSION

Rising obesity levels worldwide have placed increasing emphasis on the function of immune cells in the VAT. Here we provide evidence that during the pathogenesis of obesity, IL-10 produced by Blimp-1^+^ aTregs contributes to DIO. Under SFD conditions, we showed that Blimp-1 expression was required to support the aTreg population. Additionally, Blimp-1 maintained markers of aTreg differentiation including ST2, KLRG1 and PPAR*γ*. Under HFD conditions, Blimp-1 deficiency did not affect the frequency or number of the aTreg population in the VAT. However, with DIO loss of Blimp-1 expression in Tregs resulted in decreased IL-10 production by aTregs, increased beiging of white adipocytes in the VAT and overall less weight gain and increased protection from insulin resistance and glucose intolerance. Moreover, increasing IL-10 mRNA levels in human VAT correlated positively with increasing BMI.

Given the known function for Treg-derived IL-10 in suppressing inflammation and maintaining peripheral tolerance(*27*), we were initially surprised to observe reduced body weight and reduced insulin resistance and glucose tolerance with loss of IL-10 in the aTregs. Previous data have shown that IL-10 in the VAT comes from bone marrow-derived immune cells, which enabled us to rule out a potential role for preadipocytes, stromal cells or fibroblasts in IL-10 production(*11*). It is possible that other immune cell populations could contribute to the immune cell-adipocyte IL-10 crosstalk but as we demonstrate here, loss of Blimp-1 production by aTregs was sufficient to generate the same weight loss, insulin sensitivity and adipocyte beiging as reported with germline IL-10 deficiency(*11*).

Blimp-1 is not the only known driver of IL-10 expression in immune cells. In VAT-resident iNKT cells, referred to as NKT10 cells for their IL-10 production, it was previously shown that the transcription factor E4bp4 is a critical driver of IL-10 expression by these cells(*26*). E4bp4 is highly expressed in aTregs(*2*), suggesting that IL-10 expression in aTregs could be driven by more than Blimp-1 alone. Nevertheless, as we show here, loss of Blimp-1 expression in aTregs was sufficient to substantially reduce IL-10 production from these cells. Similarly, expression of Gata3, another transcription factor that can positively regulate IL-10 production in T cells(*28, 29*), was unaffected by loss of Blimp-1 expression in aTregs (data not shown). Although expression of the transcription factor cMaf is not overtly high in aTregs(*2*), it is increased over that of conventional Tregs and might represent another transcription factor that could regulate IL-10 production by aTregs(*30*). In T cells, IL-10 expression can be repressed by the transcription factor bHLHe40(*31, 32*). bHLHe40 is highly expressed in aTregs under standard diet conditions(*2*), indicating that an elaborate interplay between multiple transcription factors likely occurs in aTregs, enabling them to produce IL-10 in the VAT(*20*).

Finally, there is the question of timing between aTreg dysfunction and the development of obesity and its comorbidities. Previous results and our data presented here have shown that the aTreg population decreases with increasing obesity(*4*). Yet, our data also show that IL-10 produced by the residual aTreg population in obese mice helps to repress Ucp-1 expression and by extension thermogenesis of adipocytes, resulting in increased body weight and metabolic disease. We think it possible that both phenomena could occur simultaneously: increasing adiposity leading to decreased aTreg frequency, while at the same time, aTregs that remain in the VAT continue to produce IL-10 and thus suppress white adipocyte metabolic activity. Additional factors, such as changes in VAT macrophage and effector T cell populations, might also add to the systemic inflammation associated with obesity.

In humans, as previously reported(*4*), we observed that aTreg signature transcripts decrease with increasing BMI. However, we also observed that in human VAT, IL-10 transcripts increase with increasing BMI. Thus, IL-10 production continues in human VAT as adiposity increases. These data suggest that the immune cell-IL-10-adipocyte axis in the VAT might be important targets for therapeutics to treat obesity and its associated illnesses.

## MATERIALS AND METHODS

### Study design

The objective of this study was to interrogate the function of Blimp-1 in adipose resident Tregs in mice on standard chow and HFD conditions. To do this, we used a combination of in vivo and ex vivo assays on mouse samples. We designed and performed the experiments predominantly using flow cytometry and metabolic indices for our analysis. The number of replicates for each experiment is indicated in the figure legends.

### Mice

C57BL/6 mice were purchased from the Jackson Laboratory. Blimp-1-YFP (commercially available at Jackson, stock number 008828), Foxp3-RFP (commercially available at Jackson, stock number 008374)(*33*) and Blimp-1^f/f^ mice (commercially available at Jackson, stock number 008100)(*34*) were donated by A. Poholek (University of Pittsburgh). Foxp3-YFP-Cre mice were purchased from Jackson Laboratory (stock number 016959)(*35*). All experiments were approved by the University of Pittsburgh Institutional Animal Care and Use Committee. The Foxp3YFPiCre mice were purchased from Jackson Laboratories (*35*). All mice used in experiments were age-matched males between 26-28 weeks of age. Male mice were used due to the hormonal fluctuations that occur in female mice. Where indicated, mice were fed HFD (60% kcal fat, Research Diets, D12451) from 8 weeks of age until 26-28 weeks of age. Male animals were randomly assigned to groups of 3-7 mice per experiment and at least two independent experiments were performed throughout the study. Mice were bred and housed in specific pathogen-free conditions in accordance with the Institutional Animal Care and Use Guidelines of the University of Pittsburgh.

### Cell isolation

Murine gonadal adipose single-cell suspensions were prepared as previously described (*36*). Briefly, tissue was harvested, weighed, finely chopped with a razor blade and digested with Liberase TM and DNase I (Sigma-Aldrich) at 37C with shaking for 30 minutes before filtering through a 70-um nylon mesh and centrifugation. The supernatant was removed and the stromal vascular fraction (SVF) was isolated and processed for flow cytometry. Single cell suspensions of spleen cells were made by processing murine spleens between two glass slides.

### Flow cytometry

Antibodies against murine CD4 (RM4-5, AF700, BV650), CD206 (C068C2, APC), CD8 (53-6.7, APC/Fire 750, BV785), CD11c (N418, APCCy7, Pe/Cy5), CCR2 (SA203G11, BV421), CD11b (M1/70, BV421), F4/80 (BM8, BV510, BV711, FITC), TNF*α* (MP6-XT22, BV510), TCR*β* (H57-597, BV605), NK-1.1 (PK136, BV650), and PD-1 (RPM1-30, PE/Dazzle 594) were purchased from BioLegend (San Diego, CA). Antibodies against CD4 (GK1.5, BUV395), ST2 (U29-93, BV480), and GATA3 (L50-823, FITC, PE-Cy7) were purchased form BD (Franklin Lakes, NJ). PPAR*γ* antibody (polyclonal, unconjugated) was purchased from Bioss (Woburn, MA). Antibodies against B220 (RA3-6B2, AF488), CD11c (N418, e450, PE-Cy5.5), CD25 (PC61.5, PE-Cy7), CD4 (GK1.5, APC-eFluor 780, Super Bright 645), CD45 (30-F11, Pacific Orange), FOXP3 (FJK-16s, APC, e450, FITC), GITR (DTA-1, Super Bright 600), IFN*γ* (XMG1.2, PE-eFluor610), IL-10 (JES5-16E3, PE), KLRG1 (2F1, APC, APC-eFluor 780, PE-eFluor 610), NK-1.1 (PK136, PE), and ST2 (RMST2-2, PE, PerCP-eFluor710) were purchased from eBioscience (San Diego, CA). Antibodies against rabbit IgG (AF647) and viability dye (e506, near IR, UV) were purchased from Invitrogen (Carlsbad, CA).

Extracellular flow staining was performed in 2% FBS-PBS in the presence of BioLegend’s TruStain fcX (San Diego, CA) for 30min at 4C°. Fixation, permeabilization and intracellular staining were performed using eBioscience’s Foxp3/transcription factor staining buffer set per manufacturer’s directions. Flow data was collected using a Cytek Aurora or BD LSR, and analyzed using FlowJo.

### Intracellular cytokine analysis

For intracellular cytokine staining total SVF was stimulated with PMA (50ng/ml) (Sigma) and ionomycin (1nM) (Sigma) for 4-6 hours. Protein transport inhibitor (eBioscience) was added to the culture at the recommended concentration. Cells were surfaced stained, before fixing, permeabilizing and intracellular staining according to the manufacturer’s instructions (eBioscience Foxp3 staining kit).

### Determination of fasting blood glucose, insulin and glucose tolerance testing (GTT)

Mice were fasted overnight before fasting blood and insulin levels were measured. Blood glucose was measured using a handheld glucometer (OneTouch Ultra 2) and plasma insulin was measured by ELISA (Alpco). For the glucose tolerance tests (GTTs), we administered glucose (2g/kg of body weight) by intraperitoneal injection after an overnight fast. We measured changes in plasma insulin at 15, 30 and 120 minutes and blood glucose at 15, 30, 45, 60 and 120 minutes after glucose injection.

### Indirect calorimetry and body composition measurements

Indirect calorimetry was performed using the Promethion Multiplexed Metabolic Cage System (Sable Systems) by the University of Pittsburgh Center for Metabolism and Mitochondrial Medicine. Animals were placed individually in chambers for 24 hours to acclimate followed by 48 hours at ambient room temperature with 12-hour light/dark cycles for analysis. Animals had free access to food and water. Respiratory measurements (VO_2_/VCO_2_) were made at 5-minute intervals. Food intake was measured in metabolic chambers during the 48-hour period. Body composition including fat and lean mass was determined using by EchoMRI. Mice were weighted weekly to determine weight increases over time on SFD and HFD.

### Tissue hematoxylin and eosin (H&E) staining and immunohistochemistry

Tissues were placed in cassettes and submerged in 10% formalin solution overnight. Tissue cassettes were washed with tap water for 15 minutes and stored in 70% ethanol at room temperature. Paraffin embedment and sectioning (4-5 microns thickness) was performed by StageBio. Slides were deparaffinized and rehydrated using ethanol and PBS respectively. H&E staining was performed using hematoxylin for 5 minutes followed by eosin for 15 seconds. Sample slides were incubated with Ucp-1 antibody (Abcam) overnight.

### RNA sequencing and analysis

RNA was isolated from sorted Tregs (CD4^+^, CD25^+^, Foxp3-YFP^+^) from the spleen or adipose tissue. Libraries were prepared using Ultra DNA library preparation kits. RNA sequencing analysis was performed on Illumina NextSeq500 using 500bp paired-end reads by Health Science Sequencing core facility at University of Pittsburgh. Raw sequence reads were trimmed of adapter sequences using CLC Genomics Suite. The trimmed reads were mapped onto the mouse genome and gene expression values (TPM; transcripts per million kilobase) were calculated using CLC Genomics Suite Workbench 11. Differential gene expression was analyzed using Partek Genomics Suite and graphs generated using Graphpad Prism with values normalized as follows: (TPM value for gene X in condition A)/ (mean TPM of gene X in all conditions for that sample).

### Real-time PCR

RNA was extracted from epidydimal adipose tissue using TRIzol (Invitrogen), and cDNA was created using iScript cDNA synthesis kit (Biorad). TaqMan gene expression assays were performed in triplicates on the StepOnePlus PCR system (Applied Biosystems). Data were normalized to HPRT. TaqMan probes: UCP-1: Mm01244861_m1, CIDEA: Mm00432554_m1, ELOVL3: Mm00468164_m1, HPRT: Mm03024075_m1.

### Statistical analysis

All graphs were created using GraphPad Prism 8 and statistical significance was determined with the two-tailed paired or unpaired Student’s t-test or using one-way ANOVA adjusted for multiple comparisons where appropriate. For the human mRNA analysis, Spearman’s correlation was used. We used a P value of <0.05 as statistically significant.

## ACKNOWLEDGMENTS

We would like to acknowledge Drs. McGeachy and Kane for critical reading the manuscript and all members of the D’Cruz lab for their constructive criticism and comments. We would also like to thank the University of Pittsburgh Unified Flow Core for assistance with cell sorting and flow cytometry. The authors declare no competing interests.

## FUNDING

This work was supported by NIH R01 DK114012 and R01 DK119627 to M.J.J., funding to the Center for Metabolism and Mitochondrial Medicine by the Pittsburgh Foundation, and funding from the University of Pittsburgh to L.M.D.

## AUTHORS CONTRIBUTIONS

L.Y.B. conducted the experiments, analyzed the data, and wrote the manuscript; X.Q., G.J.M., A.N.F. and A.B.F. conducted the experiments; I.S., B.X. and M.J.J. performed the metabolic studies and analysis; K.E.H. and S.C.W. provided assistance with the microscopy experiments and analysis, A.C.P. provided scientific insight, L.M.D. designed the experiments, analyzed the data and wrote the paper.

## COMPETING INTERESTS

The authors declare that they have no competing interests.

## SUPPLEMENTARY MATERIALS

**Supplemental Figure 1.**
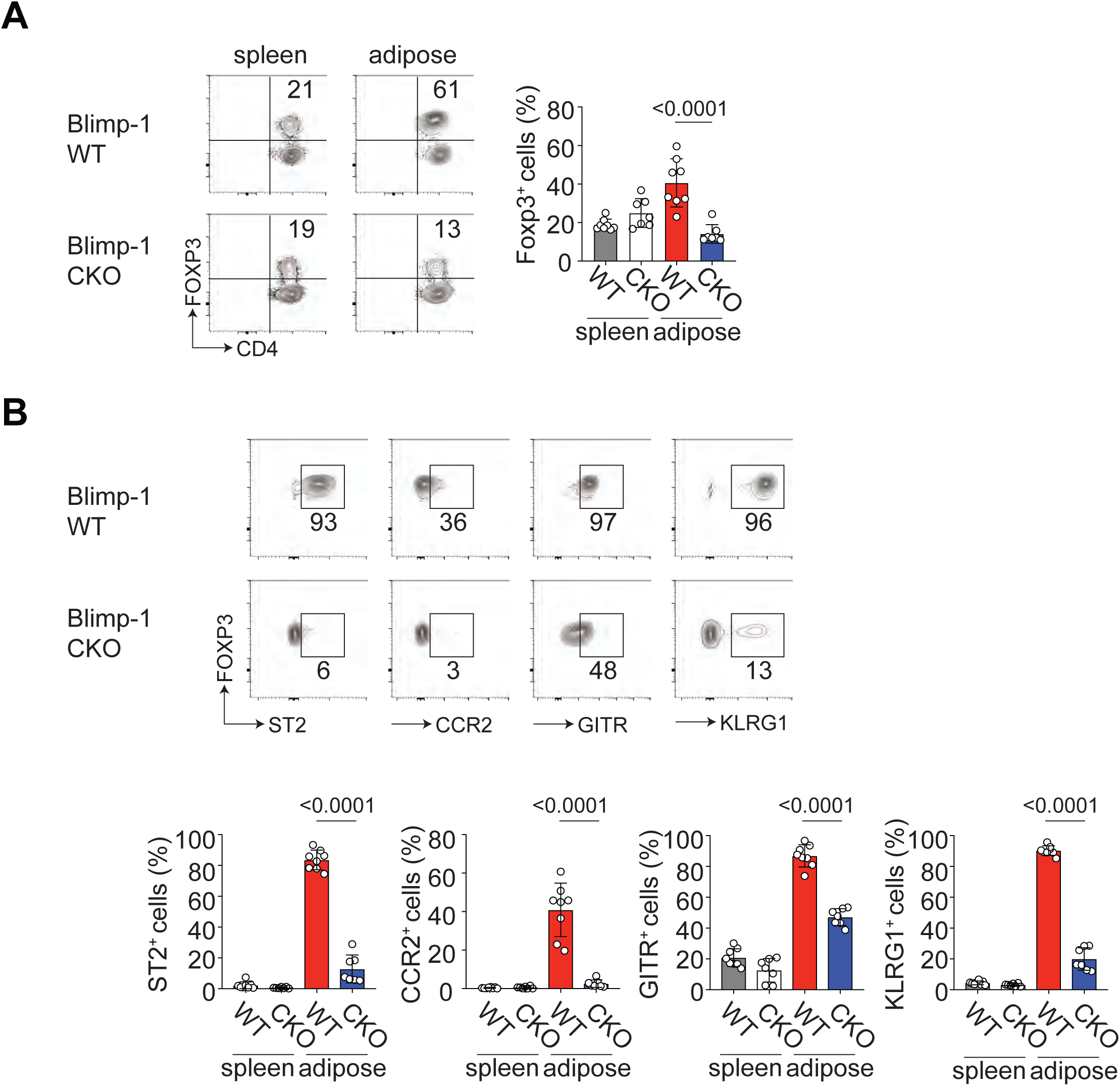
Characterization of Blimp-1-deficient Tregs from Foxp3-YFP-Cre+ conditional mice on standard fat diet (SFD). Blimp-1f/f mice were crossed to Foxp3-YFP-Cre+ mice and the Tregs from 26-28-week-old male mice on standard fat diet (SFD) were analyzed. (A) Flow cytometry plots and bar graphs showing the frequency and cell number of CD4+ Foxp3+ Tregs from the indicated tissue in Blimp-1 wildtype (WT) or Blimp-1 conditional knockout (CKO) mice. (B) Flow cytometry plots showing the frequency of gated CD4+ Foxp3+ aTregs expressing ST2, CCR2, GITR and KLRG1. Bar graphs indicate the frequency of these populations in the indicated tissues from WT or CKO mice. Data are presented as means ± s.e.m. for n = 7-8 mice per group pooled from two independent experiments. An unpaired Student’s t-test was performed to determine significance and the P values are indicated on the graphs.

**Supplemental Figure 2.**
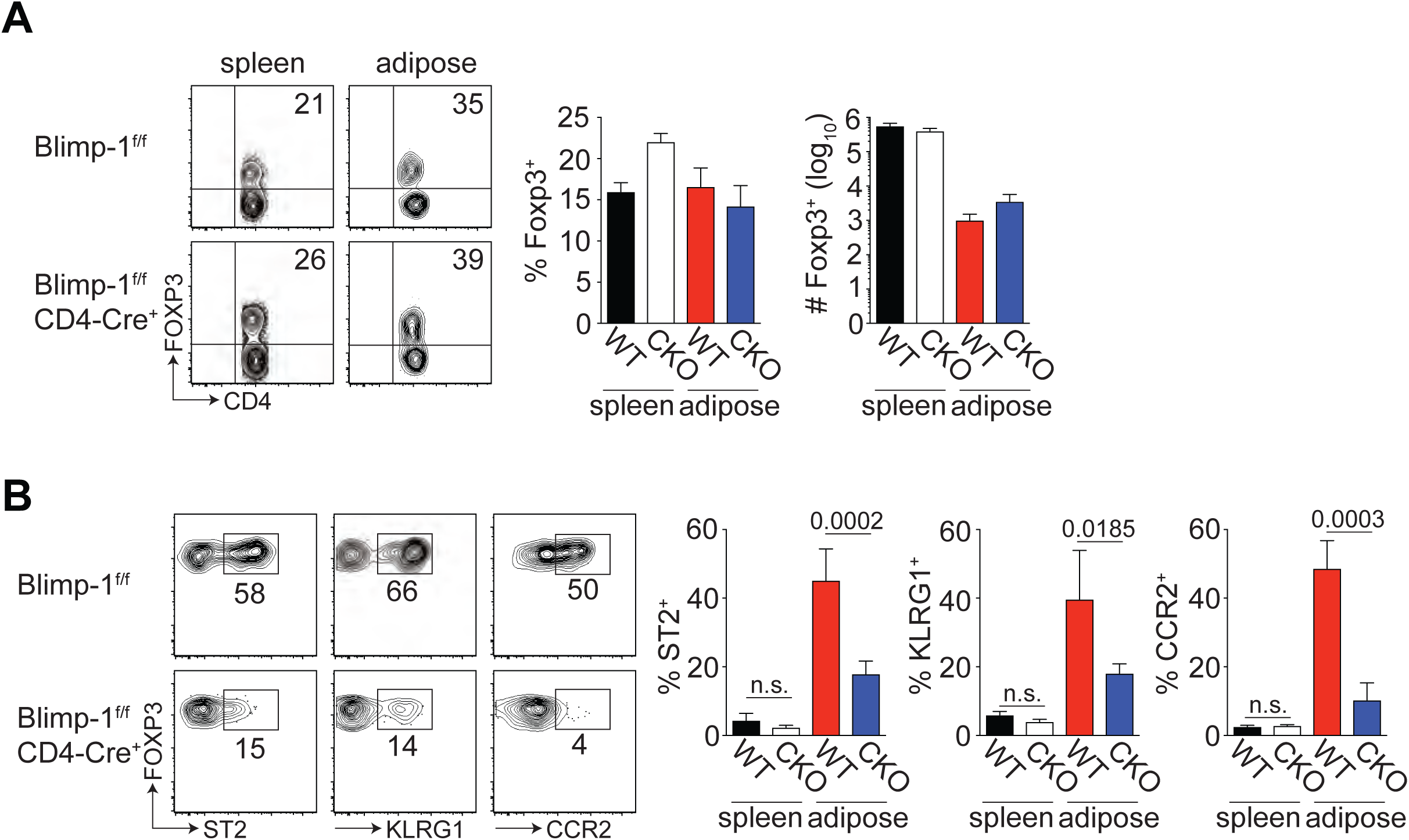
Characterization of Blimp-1-deficient Tregs from CD4-Cre+ conditional mice. Blimp-1f/f mice were crossed to CD4-Cre+ mice and the Tregs from 26-28-week-old male mice on standard fat diet (SFD) were analyzed. (A) Flow cytometry plots and bar graphs showing the frequency and cell number of CD4+ Foxp3+ Tregs from the indicated tissue in Blimp-1f/f wildtype (WT) or Blimp-1f/f CD4-Cre+ conditional knockout (CKO) mice. (B) Flow cytometry plots showing the frequency of gated CD4+ Foxp3+ aTregs expressing ST2, KLRG1 and CCR2. Bar graphs indicate the frequency of these populations in the indicated tissues from WT or CKO mice. Data are presented as means ± s.e.m. for n = 10 mice per group (A) or n = 6 mice per group (B), pooled from four independent experiments. An unpaired Student’s t-test was performed to determine significance and the P values are indicated on the graphs (n.s. = not significant).

**Supplemental Figure 3.**
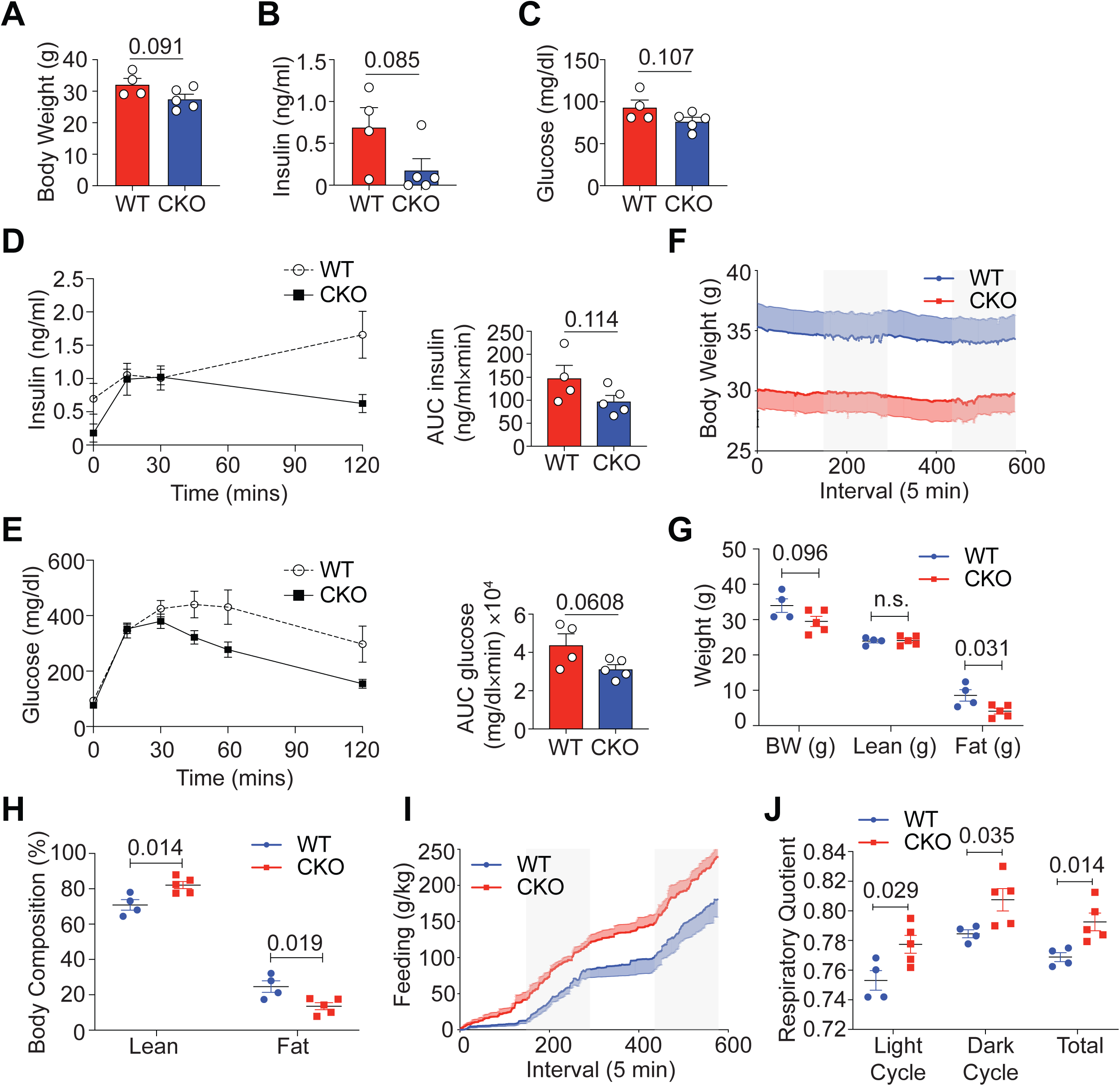
Loss of Blimp-1 expression in Tregs decreases weight and increases insulin sensitivity in mice on SFD. Male Foxp3-YFP-Cre+ (WT) and Blimp-1f/f mice crossed to Foxp3-YFP-Cre+ (conditional knockout, CKO) were placed on standard fat diet (SFD) for 18 weeks prior to metabolic analysis. (A) Bar graphs indicating body weight of 26-week old SFD-fed WT and CKO mice. (B) Bar graph showing fasting plasma insulin levels in WT and CKO mice. (C) Bar graph showing fasting blood glucose serum levels in WT and CKO mice. (D) Graph indicating serum plasma insulin levels in mice over time after i.p. glucose injection. Bar graph indicates the area under the curve (AUC) for both groups. (E) An i.p. glucose tolerance test (GTT) was performed on WT and CKO mice in (E). Bar graph indicates the area under the curve (AUC) for both groups. (F) Graph showing body weight as measured by Promethion Multiplexed Metabolic Cage System) for 48-hr total duration in WT and CKO mice. (G) Body weight (BW), lean and fat mass in grams of WT and CKO mice. (H) Percent lean and fat mass expressed per gram BW in WT and CKO mice. (I) Food intake in g/kg lean mass for WT and CKO mice. (J) Respiratory exchange ratio (RER) in light, dark and total in WT and CKO mice. For all experiments each dot represents one mouse and an unpaired Student’s t-test was performed to determine significance (A-C, F-J). Comparisons at each time point were made against WT control mice by ANOVA (D, E). Data are presented as means ⊠ s.e.m., for n = 4-5 mice per group.

**Supplemental Figure 4.**
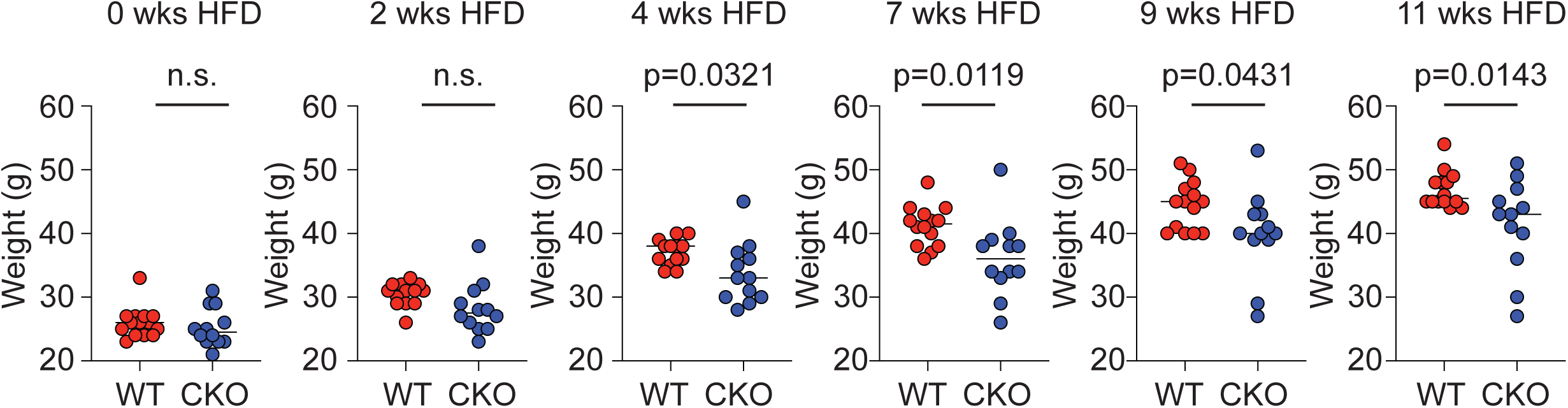
Weight gain in WT and Blimp-1-deficient Treg mice on HFD over time. Graphs showing the weight in grams of wildtype (WT) or Blimp-1 conditional knockout (CKO) mice on 60% HFD over the time indicated. Data are presented as means ± s.e.m. for n = 12 mice per group pooled from two independent experiments. An unpaired Student’s t-test was performed to determine significance and the P values are indicated on the graphs (n.s. = not significant).

**Supplemental Figure 5.**
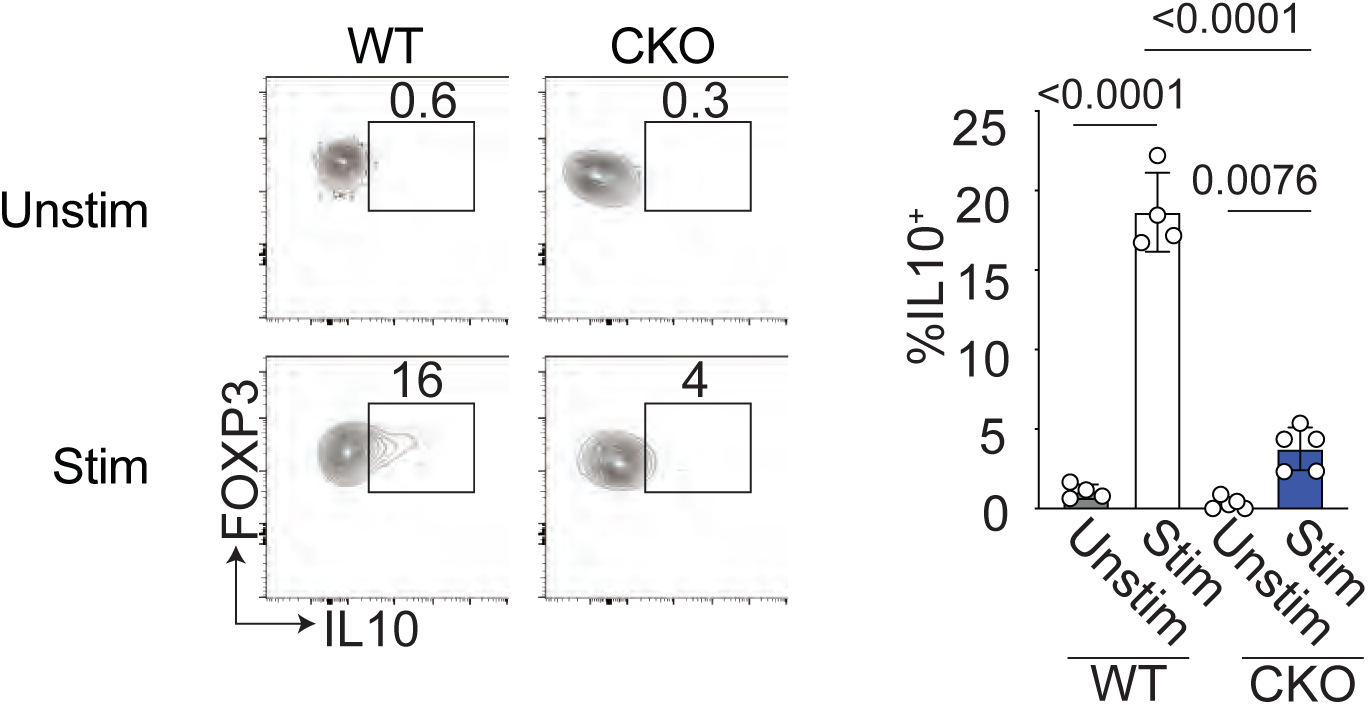
Loss of IL-10 expression by Blimp-1-deficient aTregs in SFD mice. Flow cytometry plots and bar graphs showing expression of IL-10 by gated CD4+ Foxp3+ cells that were unstimulated (unstim) or stimulated (stim) for 5 hours with PMA and ionomycin from the VAT of WT and CKO mice after 26-28 weeks on SFD. Data are presented as means ± s.e.m., for n = 4-5 mice per group. Each dot represents one mouse and an unpaired Student’s t-test was performed to determine significance.

**Supplemental Figure 6.**
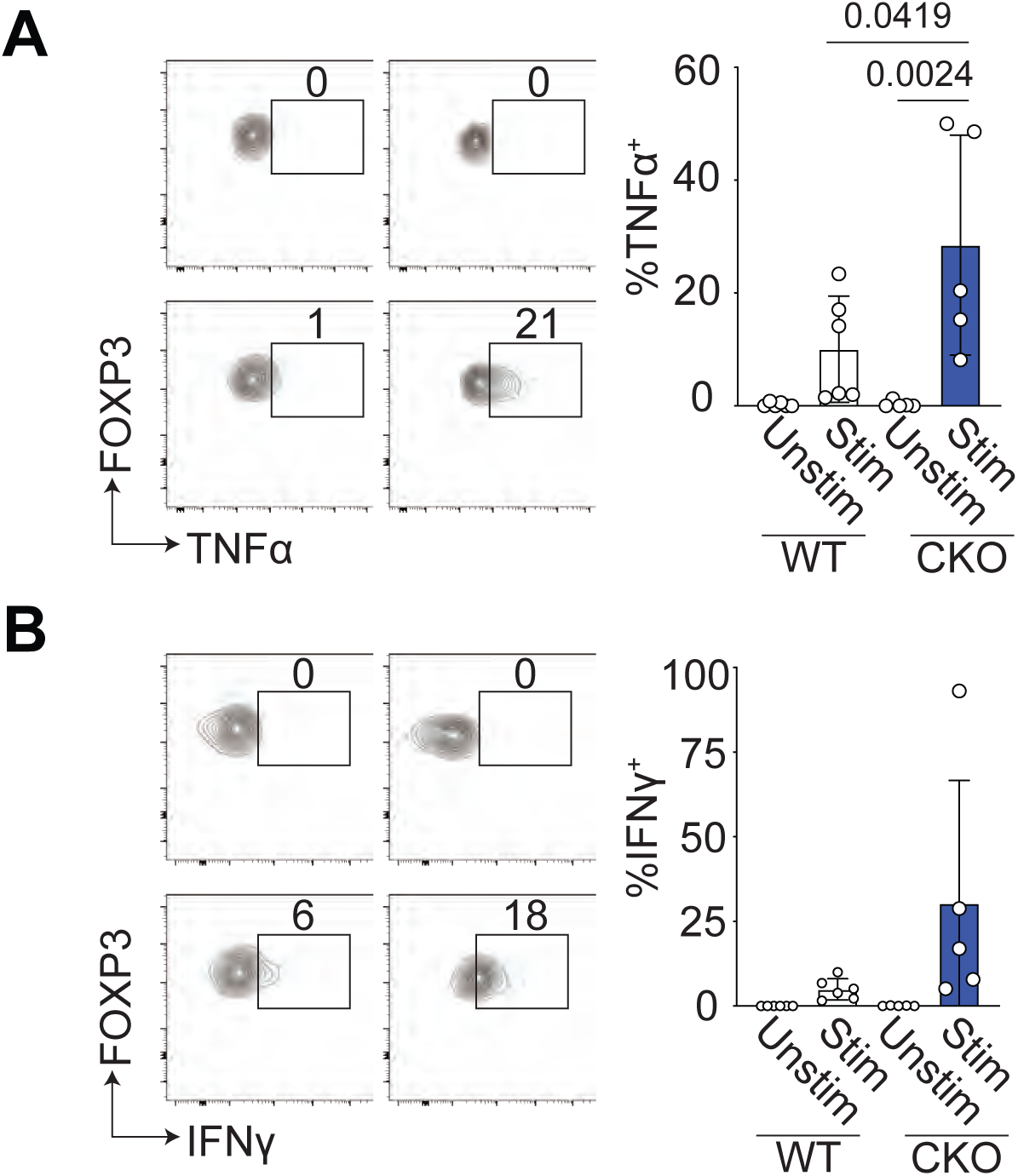
Inflammatory cytokine production by WT and Blimp-1-deficient aTregs. (A and B) Flow cytometry plots and bar graphs showing expression of TNFα and IFNγ by gated CD4+ Foxp3+ cells that were unstimulated (unstim) or stimulated (stim) for 5 hours with PMA and ionomycin from the VAT of WT and CKO mice on HFD. Data are presented as means ± s.e.m., for n =5-6 mice per group, pooled from at least 2 independent experiments. Each dot represents one mouse and one-way ANOVA was performed to determine significance.

